# Abrupt permafrost thaw triggers microbial bloom and grazer succession

**DOI:** 10.1101/2022.08.09.499897

**Authors:** Maria Scheel, Athanasios Zervas, Ruud Rijkers, Alexander Tøsdal Tveit, Flemming Ekelund, Francisco Campuzano Jiménez, Carsten Suhr Jacobsen, Torben Røjle Christensen

## Abstract

Permafrost soils store a substantial part of the global soil carbon and nitrogen. However global warming causes abrupt erosion and gradual thaw, which make these stocks vulnerable to microbial decomposition into greenhouse gases. Here, we investigated the microbial response to abrupt *in situ* permafrost thaw. We sequenced the total RNA of a 1 m deep soil core consisting of up to 26’500-year-old permafrost material from an active abrupt erosion site. We analysed the microbial community in the active layer soil, the recently thawed, and the intact permafrost and found maximum RNA:DNA ratios indicating a microbial bloom in recently thawed permafrost. Several fast-growing prokaryotic taxa dominated thawed permafrost, including Sphingobacteriales, Burkholderiales, and Nitrosomonadales. Overall, the thaw state and soil moisture consistently explained changes in community composition, with especially the permafrost community being significantly distinct from thawed soils. Predation correlated with changes in prokaryotic composition. Bacterial grazers were dominated by Myxococcales and abundant in the active layer. In contrast, protozoa, especially Cercozoa and Ciliophora, doubled in relative abundance in thawed layers. Our findings highlight the ecological importance of a rapid development of microbial blooms as well as the successive predation as biological control mechanism in abruptly thawing permafrost.

**One sentence summary:** Using total RNA from an up to 26’500-year-old abruptly eroding permafrost site in Greenland, we described a microbial bloom and its controls, including bacterial and microeukaryotic predators.

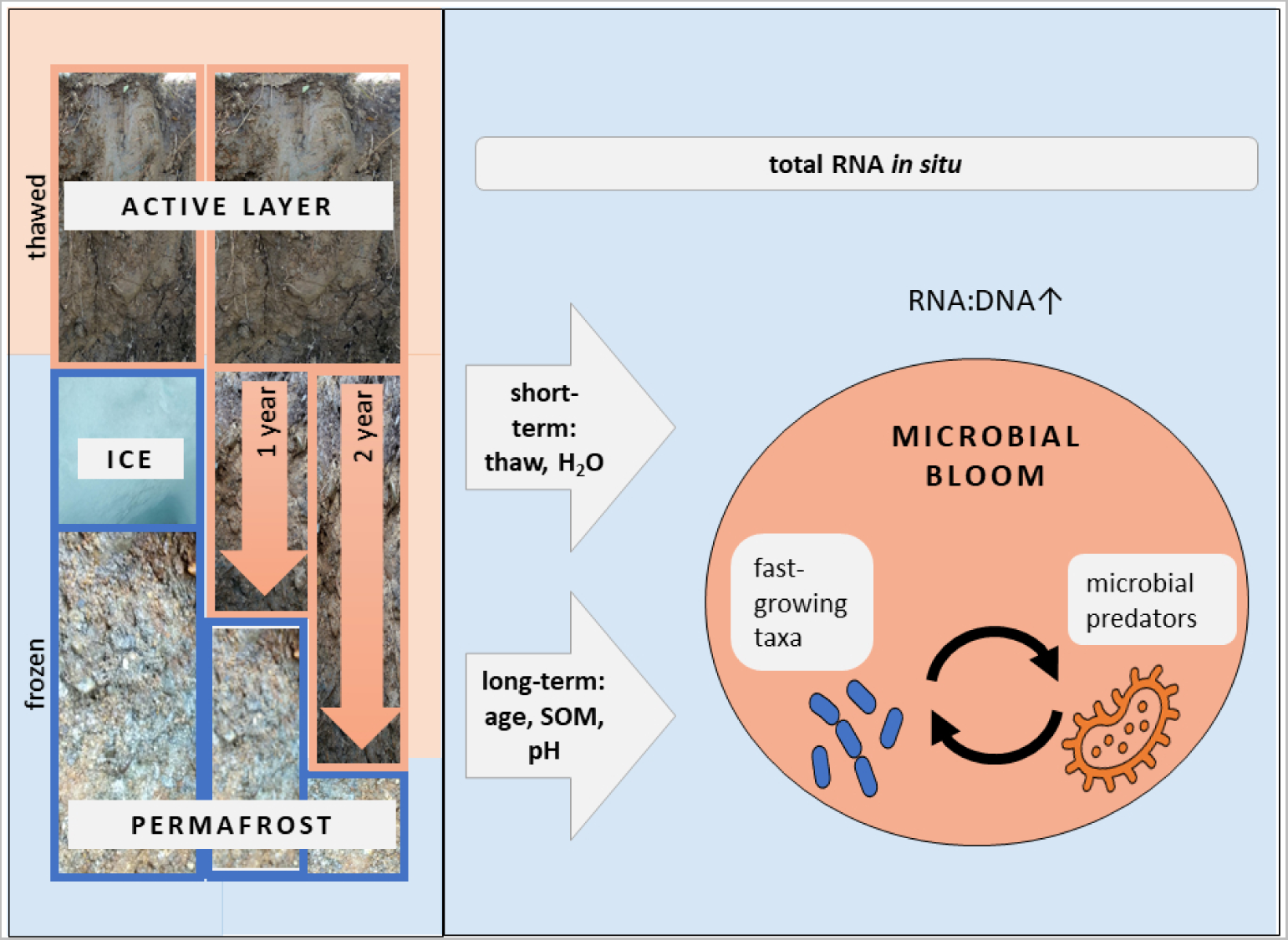

## 1. Introduction

Permafrost soils are among the most vulnerable ecosystems of the globe, both in terms of biodiversity and potential degradation in response to global warming (Abbott *et al*., 2022). Permafrost is found in historically cold regions and remains frozen for at least two consecutive years. It consists of a seasonally thawed active layer and permafrost below, often remaining frozen for millennia. These conditions selected for a unique but still elusive microbial communities under constant freezing conditions (Jansson & Tas, 2014). Due to the slow microbial decomposition at low temperatures, permafrost soils contain 1100–1500 Pg C, over half of all global soil carbon (C) (Tarnocai *et al*., 2009, Hugelius *et al*., 2014), with approximately 800 Pg C stored in permafrost (Hugelius *et al*., 2014).

Ambient temperatures in the Arctic increased up to four times faster than the global average (Rantanen *et al*., 2022) and lead to gradual thaw of permafrost, which results in annually deeper thawing active layers (AMAP, 2021). This enables a deeper rooting vegetation and potential microbial remineralisation of permafrost carbon (Schuur & Mack, 2018). However, warming also causes ground ice to melt and soils to collapse abruptly, modelled to affect up to half of all permafrost carbon by 2100 (Turetsky *et al*., 2020). However, predictions of future carbon and nitrogen release as carbon dioxide (CO_2_), methane (CH_4_), and nitrous oxide (N_2_O) from permafrost are uncertain due to high spatial variability and nonlinearity of the ecological response to permafrost warming and thaw (Monteux *et al*., 2020, IPCC, 2021, Ernakovich *et al*., 2022). Permafrost – due to long-term stable freezing temperatures – harbours liquid water only in salty brine channels and stores most of the nutrient-poor carbon as highly recalcitrant organic matter (Parmentier *et al*., 2017). These conditions select for resistant taxa that exhibit slow growth rates, higher metabolic versatility, and an ability to perform syntropy (Allison & Martiny, 2008, Shade *et al*., 2012), as observed in most anoxic systems. Permafrost communities are functionally constrained due to environmental limitations and slow reproduction rates (Monteux *et al*., 2020). The microbiome of frozen permafrost differs significantly from overlaying thawed soils, as seasonally fluctuating temperature has selected for highly resilient taxa with high growth rates (Allison & Martiny, 2008, Bardgett & Caruso, 2020). However, upon thaw, permafrost soil communities, similar to temperate soil microbiomes, often become functionally redundant (Nannipieri *et al*., 2017), both during *in situ* (Monteux *et al*., 2018) and experimental thaw (Monteux *et al*., 2020). This release of limitation was achieved through an incubation with grassland and active layer soil material transplanted in permafrost soil and during thaw resulted in higher nitrogen (N) mineralisation and release of CO_2_ (Monteux *et al*., 2020).

Other possible microbial responses to thaw include the revival of former resting stages, such as cysts and endospores (Lennon & Jones, 2011), which upon suitable thermal and moisture conditions can lead to local microbial blooms. Moreover, the active layer microbiome can colonise thawed layers following cryoturbation (Gittel *et al*., 2014), active layer detachments (Inglese *et al*., 2018), or increased root growth (Monteux *et al*., 2020). These migrations, or coalescence effects, through rapid population growth of copiotrophic taxa, can increase carbon and nitrogen release from thawed permafrost soil (Monteux *et al*., 2020).

While so far not described in thawing permafrost soils, the complexity of trophic relations play a key role in temperate soil carbon mineralisation (Bardgett & van der Putten, 2014). Particularly the predation on bacteria can act as control and driver of the soil microbiome (Thakur & Geisen, 2019, Geisen *et al*., 2021). Organisms feeding on bacteria, called bacterivores or grazers, include predatory bacteria and eukaryotes, such as protozoa (*sensu* single cell predatory protists, Geisen *et al*., 2018), nematodes, rotifers, and tardigrades (Coleman & Wall, 2015). Moreover, the selective removal of specific species and/or senescent taxa by grazers controls bacterial turnover and community composition (Trap *et al*., 2016). Lysis of prey cells and incomplete mineralisation of organic C and N by protozoa enhance the recycling and distribution of nutrients, including plant-promoting substances (Bonkowski, 2004). Although microbial predator presence, such as Myxococcales, was documented before in permafrost (Malard & Pearce, 2018, Schostag *et al*., 2019, Scheel *et al*., 2022), their relevance in Arctic soils is still illusive.

Soil microbial biodiversity highly influences the microbiome’s resilience toward climate extremes (Griffiths & Philippot, 2013, Bardgett & van der Putten, 2014) and the decrease of soil biodiversity in temperate systems was linked to increased losses of carbon to the atmosphere (Bardgett & van der Putten, 2014, Anthony *et al*., 2020). Eukaryotic microbial diversity is particularly understudied due to biases and difficulties related to the use of gene-specific primers (Harder *et al*., 2016), but the use of ribosomal RNA (rRNA) community analysis enables a less biased understanding of the putatively active total community.

Within the Arctic, particularly Greenland and abrupt erosions sites are highly understudied (Malard & Pearce, 2018, Metcalfe *et al*., 2018). Here, we overcame these limitations by sampling an *in situ* abrupt erosion site of Greenlandic permafrost and performing the first Total RNA metatranscriptomic analysis of the entire active microbial community in such an environment.

We tested if trophic dynamics impact the microbial community more than abiotic soil properties in different stages of thaw. We furthermore hypothesised that the freshly thawed soils support copiotrophic taxa that either invaded from the active layer or consist of reactivated permafrost resting stages. These insights can help us understand the spatiotemporal and trophic dynamics of permafrost microbial ecology and estimate the ecological response of gradual and rapid erosion as a potential key driver of both permafrost carbon vulnerability.

## 2. Materials and Methods

### 2.1. Soil sampling

Sampling took place in 2020 in Zackenberg valley, NE Greenland (74°30’N, 20°30’W, Fig. 1 A-B). This wide lowland valley is dominated by continuous permafrost and a vegetation of wet hummocky fens, low shrub, and graminoids (Elberling *et al*., 2008). Average temperatures varied between −2 °C in summer and −14 °C in winter between 1997 and 2006 (Christiansen *et al*., 2008) The active layer seasonally thaws between 40 cm and 2 m deep, but an increase of 0.77 cm per year when including data from 1995–2020 (Westermann *et al*., 2015, Westergaard-Nielsen *et al*., 2018).

**Fig. 1.**
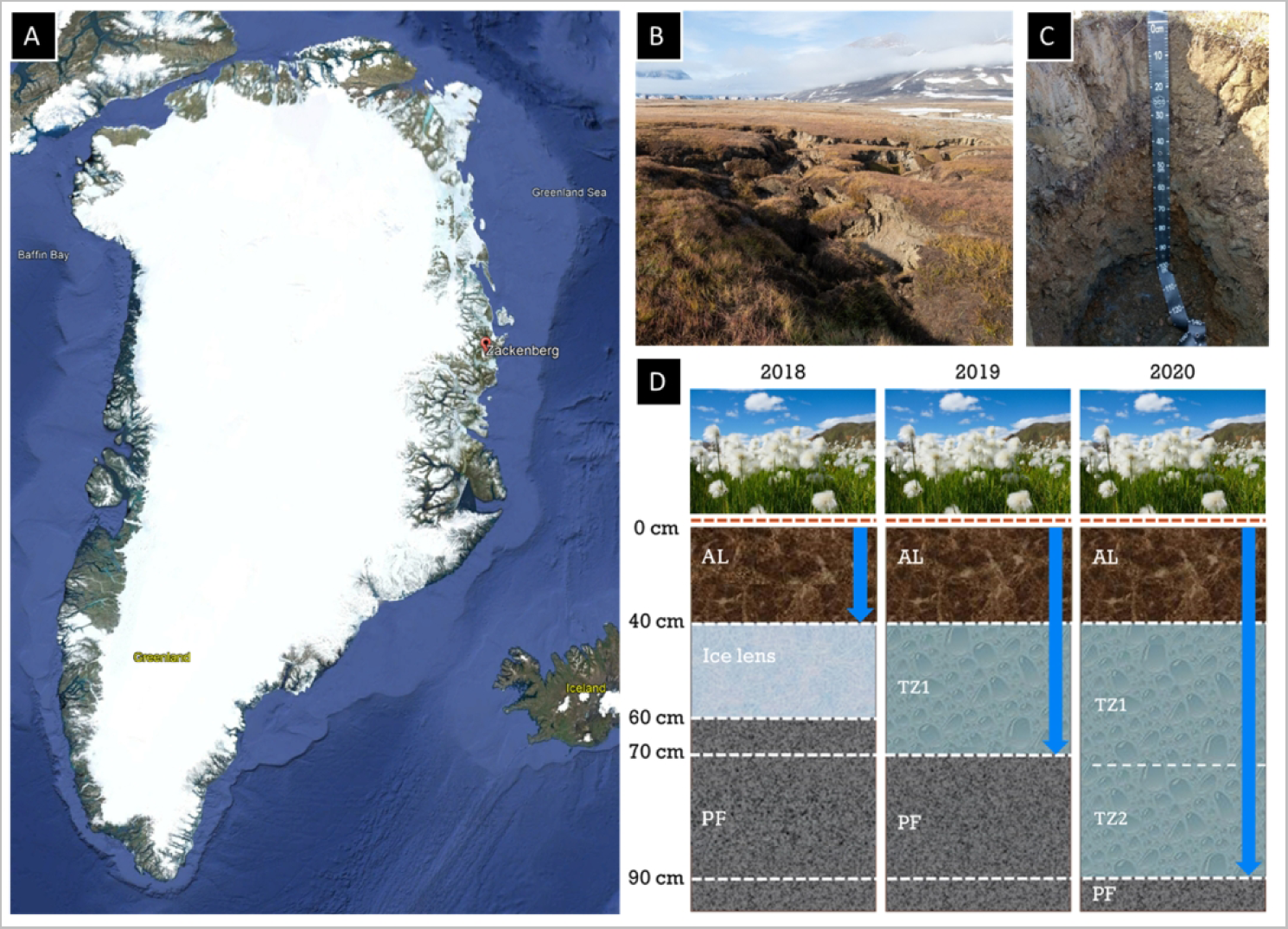
(A) Sampling site Zackenberg in Northeast Greenland, credit: Google Earth. (B) Sampling site in 2018 after initial permafrost collapse. (C) Soil profile from the surface until still frozen depth at 90 cm during sampling in 2020. (D) Scheme of abrupt permafrost thaw (indicated with blue arrows), depicting the soil profile until the permafrost (PF) layer at 90−100 cm depth from the moment of collapse in 2018 with visible ice lens below long-term active layer (AL) to the formation of transition zones (TZ, = thawed permafrost). Soil samples were taken in 2020.

The permafrost soil surface collapse abruptly in 2018 as a formerly described recent, thermal erosion gully developed in the vicinity of the Zackenberg Research Station (Christensen *et al*., 2020, Scheller *et al*., 2021, Scheel *et al*., 2022). Below the active layer (AL) at 40 cm depth, an ice lens had melted in 2019, creating a first transition zone depth until 70 cm depth (TZ1). In 2020 thaw deepened until 90 cm depth in 2020 (TZ2), while below 90 cm depth intact permafrost (PF) persisted (Fig. 1 C-D). In 2020, three replicate soil samples were taken aseptically per 10 cm intervals until a depth of 1 m, resulting in 30 samples. Due to different RNA stability at varying freezing temperatures (Schostag *et al*., 2020) and laboratory limitations in the sampling station, the samples were stored at −20 °C until transported frozen to Denmark, where they were stored at −80 °C.

### 2.2. Physicochemical soil analysis

Physical soil properties were determined in technical triplicate as described in (Scheel *et al*., 2022) from thawed 10-cm soil horizon samples until 100 cm depth. First after air-drying at 70 °C for 48 h, followed by burning at 450 °C for 2 h, the samples were weighed to determine the loss of relative weight-based soil water (H_2_O) and organic carbon (SOM, (Wilke, 2005). The pH was measured after adding 50 ml of 1 M KCl to 10 ml of airdried soil samples with a Mettler Toledo FiveEasy PlusTM pH Meter (Mettler Toledo GmbH, Gieben, Germany).

Radiocarbon dating was performed per 10-cm horizon by sifting thawed soil with a 0.5-mm sieve retaining macro plant residues and excluding roots. Per depth, triplicates were pooled, treated with HCl and NaOH, graphitised, and ^14^C isotope activity was measured using an accelerator mass spectrometer (Radiocarbon Dating Laboratory, Lund University, Lund, Sweden). The resulting age was calibrated with IntCal13 (Reimer *et al*., 2016) to ^14^C years in BP (before present = AD 1950) and the Levin post-Bomb calibration (Levin & Kromer, 2016) for results in fM (fraction modern) after 1963.

### 2.3. Nucleic acid co-extraction, library preparation, and sequencing

Deep-frozen samples were homogenised in antiseptic mortars. Then we co-extracted the total RNA and DNA of the biological replicates on up to 0.35 g frozen soil sample with the NucleoBond RNA Soil Mini kit (Macherey-Nagel GmbH & Co. KG Dueren, Germany) according to the manufacturer’s protocol. We used G2 DNA/RNA Enhancer infused 1,4 mm beads (Ampliqon, Odense, Denmark) instead of the ones provided with the kit. To remove potential DNA, the RNA extracts were treated with the DNase Max Kit (QIAGEN), following the manufacturer’s protocol. Both the removal of DNA and final RNA concentrations were evaluated with were confirmed using a Qubit® 4 Fluorometer (Thermo Fisher Scientific, Life Technologies, Roskilde, Denmark). We quality-assessed the final extracts with a TapeStation 4150 (Agilent Technologies, Santa Clara, California, US) using a high sensitivity assay. Resulting RIN values were low (1.0 ± 0.6, Supp. Tab. 1) and samples from 80–100 cm depth had RNA concentrations below detection limit. Nucleic acid concentrations were normalised for sample weight and used for extraction ratios of extracted RNA to DNA (RNA:DNA, Tab 1). We fragmented the RNA, synthesised cDNA, and metabarcoded it, using the NEBNext Ultra II Directional RNA Library Prep Kit and the NEBNext Multiplex Oligos for Illumina (New England BioLabs, Ipswich, MA, USA), following the manufacturer’s protocol. The resulting samples were pooled into equimolar metatranscriptome libraries to secure even sequencing coverage. We performed the sequencing in-house (Department of Environmental Science, Aarhus University, Denmark) on an Illumina NextSeq 500 with a v2.5 high-throughput 300 cycles kit (both Illumina, San Diego, CA, United States).

### 2.4. Bioinformatic Processing

The 352 Mio. raw paired-end Illumina reads (SRA accession number: PRJNA939404, Tab. 1) were quality-controlled with TrimGalore (https://www.bioinformatics.babraham.ac.uk/projects/trim_galore/) and filtered by removing adapters and short reads (<60 nt). These sequences were then sorted using SortMeRNA (Kopylova *et al*., 2012) into small subunit (SSU) rRNA, large subunit (LSU) rRNA, and non-rRNA sequences. The SSU reads were assembled into full-length SSU rRNA contigs with MetaRib (Xue *et al*., 2020). These contigs were taxonomically classified as described by Anwar and colleagues (Anwar *et al*., 2019), using CREST (Lanzen *et al*., 2012) against the SILVA database v.138 (Quast *et al*., 2013). The rRNA reads were mapped to resulting EMIRGE contigs using BWA (Li & Durbin, 2009), resulting in taxonomically annotated rRNA contig abundance across the 30 samples. The automatic pipeline used is available (Campuzano Jiménez, 2023).

**Tab. 1.** Soil properties include ^14^C (radiocarbon) dating results (* = fM; **BP), relative weight-based soil moisture (H_2_O), relative weight-based soil organic matter content (SOM), pH and layer, as defined by thawing processes as active layer (AL); deepest thaw in 2019: transition zone 1 (TZ1) and 2020: TZ2; and permafrost (PF). Extraction and sequencing output included the calculated RNA:DNA ratio based on co-extraction of nucleic acids in ng yield per gram dry weight (gDW). The cumulative sum of clean total RNA sequencing reads per depth. Standard deviations for triplicates per depth are indicated where available (±).

| depth [cm] | <sup>14</sup>C [*fM **BP] | H<sub>2</sub>O [%] | SOM [%] | pH | layer | DNA ng/gDW | RNA ng/gDW | RNA:DNA | total reads |
| --- | --- | --- | --- | --- | --- | --- | --- | --- | --- |
| 0-10 | 1.04* | 28.80 ± 3.38 | 8.73 ± 1.82 | 4.22 ± 0.03 | AL | 835.55 ± 391.99 | 357.46 ± 171.55 | 0.46 ± 0.16 | 1.25E+07 |
| 10-20 | 1.13* | 22.52 ± 0.73 | 5.39 ± 0.83 | 4.02 ± 0.05 |  | 82.65 ± 93.13 | 72.99 ± 8.38 | 1.78 ± 1.28 | 4.85E+07 |
| 20-30 | 1.16* | 22.05 ± 3.30 | 10.19 ± 3.22 | 4.29 ± 0.01 |  | 561.38 ± 571.54 | 91.19 ± 1.99 | 0.29 ± 0.19 | 2.25E+07 |
| 30-40 | 1.2* | 26.57 ± 0.83 | 13.68 ± 0.44 | 4.25 ± 0.01 |  | 186.67 ± 33.49 | 86.43 ± 23.06 | 0.49 ± 0.20 | 6.07E+07 |
| 40-50 | 2 635** | 7.73 ± 1.58 | 2.59 ± 0.68 | 4.63 ± 0.03 | TZ1 | 54.68 ± 60.24 | 40.66 ± 42.59 | 0.83 ± 0.91 | 2.39E+07 |
| 50-60 | 3 770** | 15.61 ± 6.28 | 2.93 ± 0.30 | 4.13 ± 0.02 |  | 10.99 ± 9.45 | 40.51 ± 10.58 | 9.34 ± 11.03 | 7.22E+07 |
| 60-70 | 26 500** | 8.72 ± 2.53 | 1.52 ± 0.16 | 4.86 ± 0.05 |  | 8.66 ± 11.41 | 36.19 ± 4.39 | 11.98 ± 9.47 | 4.90E+07 |
| 70-80 | 22 100** | 6.35 ± 0.60 | 1.10 ± 0.24 | 4.48 ± 0 | TZ2 | 2.11 ± 11.97 | 17.94 ± 31.07 | 18.48 ± 32.02 | 1.49E+07 |
| 80-90 | 26 200** | 7.96 ± 0.61 | 1.00 ± 0.20 | 4.57 ± 0.01 |  | 1.04 ± 1.80 | NA | NA | 3.46E+07 |
| 90-100 | 26 200** | 7.80 ± 0.59 | 1.04 ± 0.26 | 4.91 ± 0.01 | PF | 0.96 ± 1.66 | NA | NA | 1.40E+07 |

### 2.4. rRNA processing

The taxonomic sequence abundance was scanned for previously documented protozoan taxa: Amoebozoa, Cercozoa, Ciliophora, Euglenozoa, Foraminifera, and Heterolobosa (Geisen *et al*., 2015). Bacterial predators included Myxococcales (excluding *Sorangium*), Bdellovibrionales, *Lysobacter, Daptobacter*, and *Vampirococcus* (Petters *et al*., 2021). The rRNA relative abundance was calculated and collapsed taxonomically into prokaryotes, eukaryotes, and further into bacterivorous bacteria and protozoa.

### 2.5. Statistical analysis and data processing

We performed community analysis with the R software v.4.2.2 in R studio (R Core Team, 2022; R Studio Team, 2022), using the phyloseq (McMurdie & Holmes, 2013) and vegan packages (Oksanen *et al*., 2020). We excluded the triplicate 1 of 30 (for the depth 0–10cm) for all downstream analysis, due to significantly low number of reads (360 opposed to an average 12 Mio. reads per triplicate). Shannon (S) alpha diversity was calculated on the total number of rRNA contigs and abundance of sequence reads mapped per sample (Supp. Tab. 1). The significance of environmental data (age, pH, SOM, H_2_O, and layer) and of bacterial and protozoan predator abundance were tested with the *anova* function (PERMANOVA, 999 permutations) on Bray-Curtis dissimilarities of relative abundance per taxon within the sample to account for different read coverage across samples. For predation effects, we performed the tests on Bray-Curtis dissimilarities of only non-predatory prokaryotic and eukaryotic communities. We performed consecutive Tukey’s HSD post hoc tests to determine significant difference between taxonomic groups with layer and age as explanatory variables. Variance partitioning was performed on PERMANOVA results for thaw-related (layer and H_2_O), long-term soil (SOM, pH, age) as well as biotic (bacterial and protozoan predation) parameters. Non-metric multidimensional scaling (NMDS) plots were ordinated graphically using Bray-Curtis dissimilarities between samples of the subset communities.

## 3. Results

### 3.1. Physicochemical soil properties

Radiocarbon results differed greatly with depth, indicating three age categories, that were used for downstream statistical analysis. A more recent upper consisted of organic material and silt (AY), a deeper 2’635−3’770-year-old inorganic silt layer (AM), and the underlying deepest 22’100−26’500-year-old layer of inorganic sand and gravel (AO, Tab. 1). Soil moisture was highest in the active layer until 40 cm depth and then stayed rather stable at 6.35−8.72%, while pH stayed stable throughout between 4.02 and 4.91 (Tab. 1). Despite low biomass (174.4 ± 334.7ng/gDW DNA; 47.3 ± 111.7 ng/gDW RNA), extraction of RNA and DNA was successful in all triplicates and RNA:DNA ratios indicated higher values in the transition zones (Supp. Fig. 1, Supp. Tab. 1).

### 3.2. Diversity, ordination, and drivers of variance in community composition

The overall community composition significantly differed with depth (PERMANOVA, F.Model = 3.95, R2 = 0.331, p = 0.001), and with layer (PERMANOVA, F.Model = 3.450, R2 = 0.301, p = 0.005). Herein, especially active layer and permafrost differed significantly for the overall (Tukey HSD test, adj. p = 0.035) as well as the prokaryote (Tukey HSD test, adj. p = 0.039) communities. Furthermore, both SOM (PERMANOVA, F.Model = 2.97, R2 = 0.271, p = 0.011) and H_2_O (PERMANOVA, F.Model = 3.22, R2 = 0.287, p = 0.010) explained a significant proportion of the variation both overall and within the prokaryotic communities. Age and pH did not correlate with the total or any partial community composition (prokaryotes, eukaryotes, bacterial and eukaryotic bacterivores; Tab. 2). All sub-communities except bacterial predators were significantly impacted by layer, indicating a significantly distinct composition in AL and PF, based on Tukey HSD pairwise comparisons, although only for eukaryotes, the PF composition was significantly different from all other layers (p < 0.03, Tab. 2). Non-predatory community composition also significantly correlated with changes in SOM and H_2_O. We found strong correlation among abiotic and biotic soil parameters (Supp. Tab. 2). Especially soil moisture, organic matter content and layer significantly correlated (ANOVA, p <0.005). Variance partitioning indicated that a combination of thaw-related (layer, soil moisture), long-term soil properties (age, SOM, pH) and biotic properties (bacterial and eukaryotic predator abundance) together only explained 36.1%, and each of these factors alone only accounted for 9.1%, 6.5% and 2.2% of the variance in beta diversity of the whole community, respectively (Fig. 2).The ordination of subset communities indicated a gradual transition from AL to TZ1 and TZ2 samples (Fig. 3E-H), confirming the PERANOVA test. Taxonomic richness for annotated rRNA contigs on average reached 3652 ± 231, while Shannon (S) alpha diversity was on average 6.58 ± 0.62 for the total community and reached a maximum at 40 cm depth (Supp. Tab. 1). Prokaryotes reached the highest diversity with S > 7 in the active layer (Fig. 3I), although lower for bacterial predators and therein decreasing within the active layer (Fig. 3J). Eukaryotic and thereby also protozoan alpha diversity on average was higher in the active layer than in deeper soil (Fig. 3K-L).

**Fig. 2.**
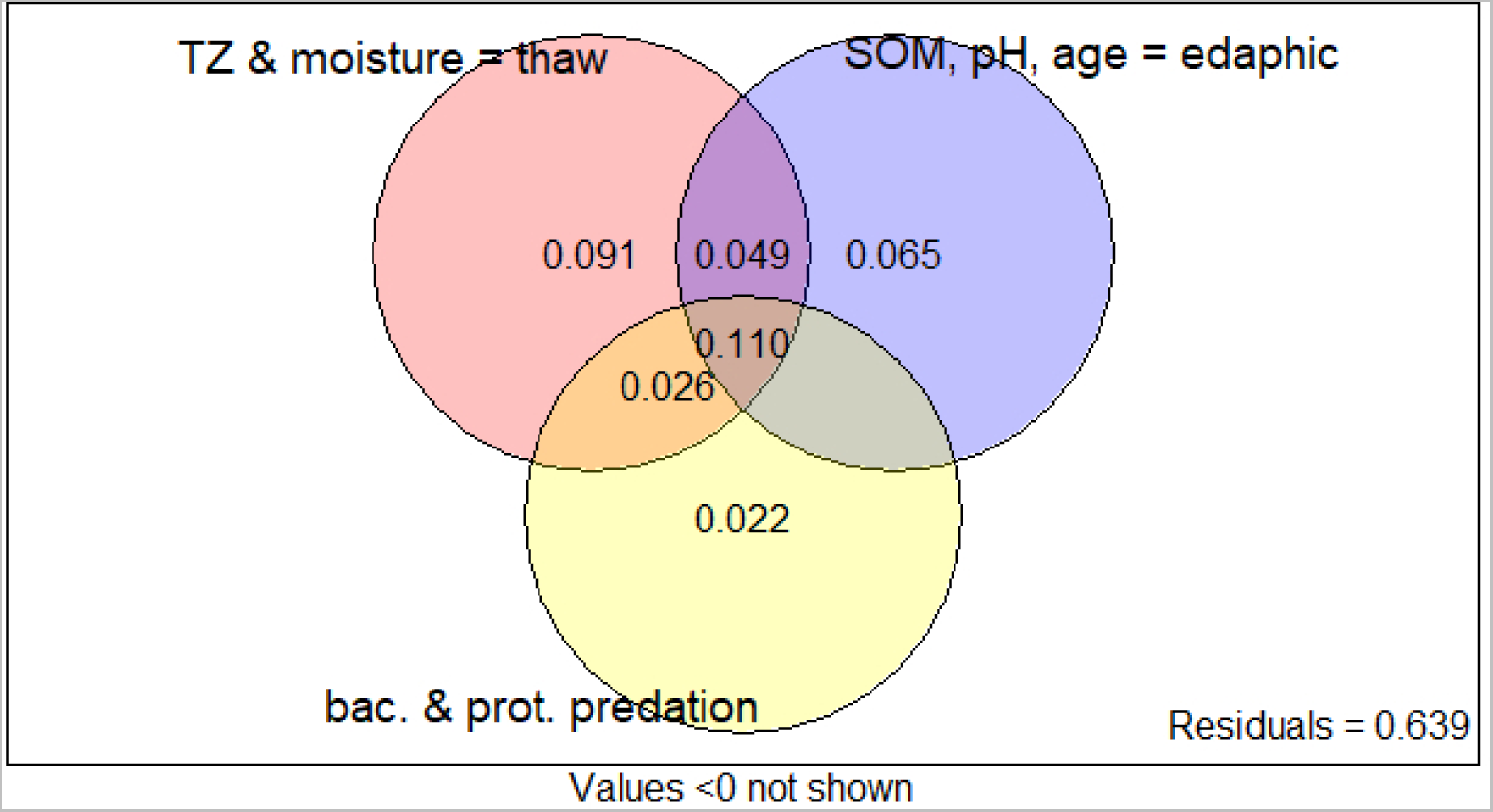
Venn diagram illustrating variance partitioning which was performed on thaw-related (layer, H_2_O), long-term soil (SOM, pH, age) as well as biotic parameters (bacterial and protozoan predator abundance) and their correlation with variance in the overall community composition. Values are given for R_2_ values, based on PERMANOVA results (999 permutations) on Bray-Curtis dissimilarities.

**Fig. 3.**
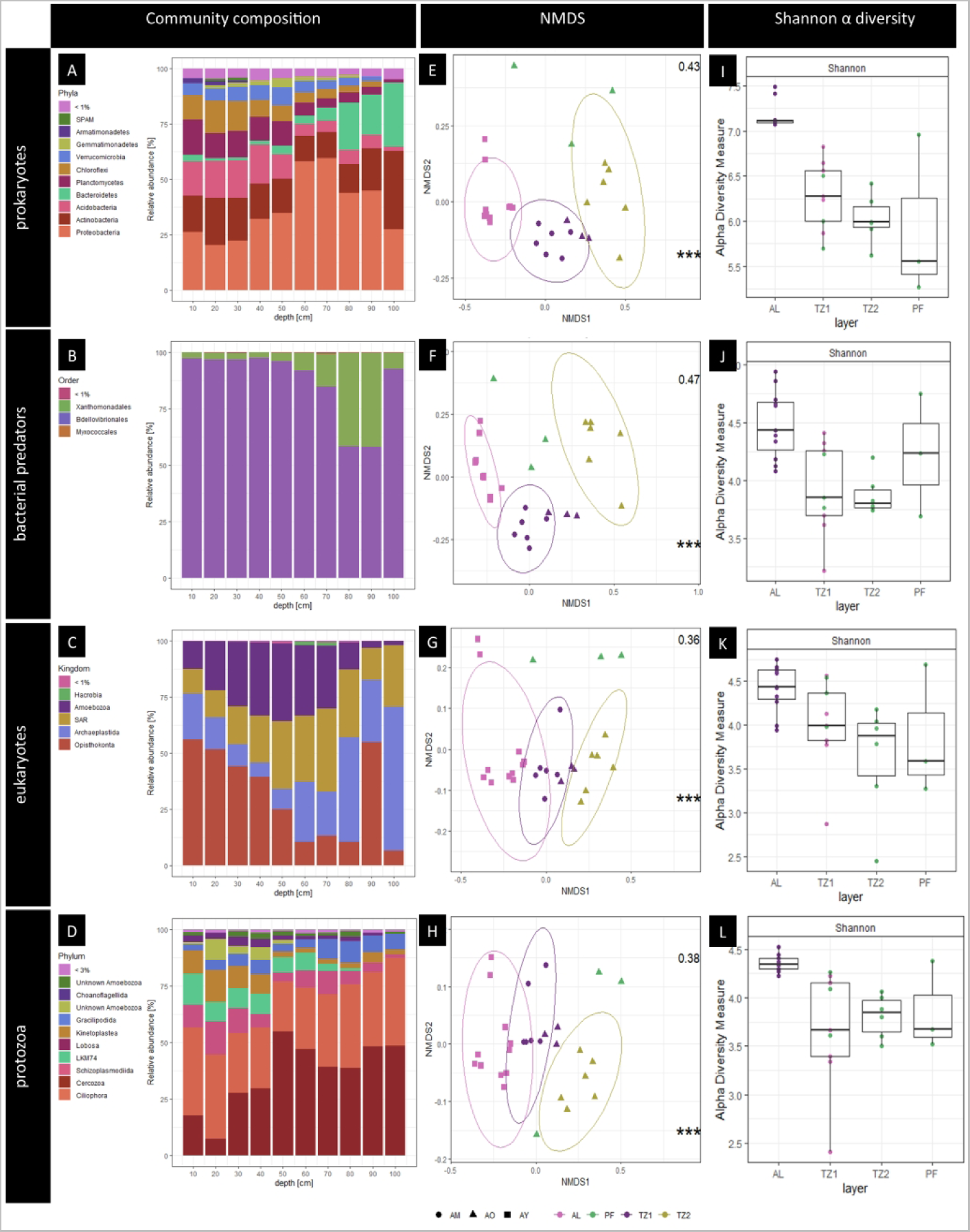
All figures are given for subsets of the total community with prokaryotes (top row), bacterial predators (second row), eukaryotes (third row) and protozoan rRNA (last row). Bar plots indicating the relative mean abundance per depth (right column) for varying taxonomic orders. Non-metric multidimensional scaling (NMDS, middle column) ordination plots performed on rRNA contig abundances per sample include R_2_ value (top right corner) and indication of significant difference across layers with p <0.05 = *, p<0.01 = **, p<0.005 = ***. Colours indicate different layers and shape different age horizons. Environmental parameters here are given as layer (active layer AL, transition zone 1 TZ1 and 2 TZ2, and permafrost PF) as well as age with young soils (AY), 2 634−3 770-year-old soil of medium age (AM), and old material (AO) of up to 22 100−26 500 years ago. Shannon alpha diversity per layer is given as boxplots with whiskers indicating standard deviation across triplicates.

**Tab. 2.** Results of permutational multivariate analysis on the mean Bray-Curtis dissimilarities per depths between total community, prokaryotic, bacterial predators (bacpred), eukaryotic and protozoan rRNA contigs. These were tested against the age categories, layer, soil organic matter (SOM) and moisture (H2O), and pH. When testing the impact of bacterial predators (bacpred) and protozoa abundance on the non-predatory prokaryotic and eukaryotic community variance was used in order to prevent artifacts. P-values < 0.05 were considered significant with * p<0.05, ** p<0.01, *** p<0.005. Non-significant categories and pair-wise contrasts, such as age, AL-TZ1, AL-TZ2, TZ1-TZ2, AY-AM, AY-AO, AM-AO were removed. Extended data on predatory and potential prey abundances can be found in SI Table 2.

| env.<br>Parameter | total |  |  | prokaryotes |  |  | bacpred |  |  | eukaryotes |  |  | protozoa |  |  |
| --- | --- | --- | --- | --- | --- | --- | --- | --- | --- | --- | --- | --- | --- | --- | --- |
|  | F.Model | R<sup>2</sup> |  | F.Model | R<sup>2</sup> |  | F.Model | R<sup>2</sup> |  | F.Model | R<sup>2</sup> |  | F.Model | R<sup>2</sup> |  |
| depth | 3.95 | 0.331 | *** | 3.90 | 0.328 | *** | 3.31 | 0.292 | * | 4.89 | 0.379 | *** | 3.48 | 0.303 | *** |
| layer | 3.45 | 0.301 | ** | 3.46 | 0.302 | *** | 2.15 | 0.212 |  | 3.28 | 0.291 | * | 2.63 | 0.247 | * |
| AL-PF |  |  | * |  |  | * |  |  |  |  |  | ** |  |  | * |
| TZ1-PF |  |  |  |  |  |  |  |  |  |  |  | * |  |  | * |
| TZ2-PF |  |  |  |  |  |  |  |  |  |  |  | ** |  |  |  |
| SOM | 2.97 | 0.271 | * | 2.98 | 0.271 | * | 1.98 | 0.198 |  | 2.84 | 0.262 | * | 1.82 | 0.185 |  |
| H<sub>2</sub>O | 3.22 | 0.287 | * | 3.21 | 0.286 | * | 2.11 | 0.209 |  | 3.38 | 0.297 | ** | 2.26 | 0.220 |  |
| pH | 1.94 | 0.195 |  | 1.93 | 0.195 |  | 1.42 | 0.151 |  | 2.14 | 0.211 | * | 1.65 | 0.171 |  |
| bacpred | 2.44 | 0.234 | * | 2.32 | 0.225 | * | - | - |  | 2.22 | 0.217 |  | 2.18 | 0.214 |  |
| protozoa | 2.51 | 0.239 | * | 2.63 | 0.247 | * | 1.26 | 0.136 |  | 2.31 | 0.224 |  | - | - |  |

### 3.3. Total rRNA community composition

On average, 74.6% of all reads could be annotated as SSU rRNA and 65.5% of the trimmed reads mapped back to the assembled full-length rRNA genes. In total, 5989 full-length rRNA gene contigs were constructed with an average length of 1474 bp. Of these, only 3729 were annotated on domain level, which were utilised for all downstream analysis. Overall, 3483 rRNA contigs were assigned as Bacteria (88.9 ± 5.1%), 230 as Eukarya (10.9 ± 5.2%), and 11 as Archaea (1 ± 1%), but annotation success decreased above class level (Supp. Tab. 1). Most contigs were omnipresent. Yet, 1668 rRNA contigs were most abundant in AL (44.7% of contigs, 30.6% of all counts), while 196 rRNA contigs were most abundant specifically in TZ2 (5.3% of contigs, 8.9% of all counts) and 136 rRNA contigs had the highest relative abundance in permafrost (3.7% of contigs, 3.8% of all counts), indicating the highest number of reads per contig ratios and thus potential activity in the transition zone samples. Furthermore, the overall most abundant taxa were most abundant in the transition zones (Supp. Fig. 3).

#### 3.3.1. Prokaryotic community composition and distribution with depth

The prokaryotic community was dominated by Proteobacteria (32.5 ± 11.7% of total counts), Actinobacteria (15.6 ± 5.7% of total counts), Acidobacteria (9.6 ± 5.3% of total counts), Planctomycetes (7.3 ± 4.0% of total counts), and Chloroflexi (6.1 ± 4.3% of total counts, Fig. 3A). Further 37 bacterial and five archaeal phyla were present. These included three methanogenic rRNA contigs (0.01 ± 0.00% of total counts) and 23 methanotrophic rRNA contigs (0.97 ± 0.00% of total counts). Proteobacteria increased in relative abundance between 50−90 cm, the depth of the two thaw layers. This phylum was dominated by Betaproteobacteria (16.0 ± 15.0% of total counts) and therein by the dominant orders Nitrosomonadales (8.9 ± 10.27% of total counts) and Burkholderiales (3.2 ± 3.0% of total counts) with maximum abundances at 70 cm and 90 cm depth, respectively.

Deltaproteobacteria (9.3 ± 6.6% of total counts) decreased below 50 cm, mainly represented by Myxococcales abundances (7.1 ± 6.1% of total counts). The abundance of Actinobacteria stayed stable, while Acidobacteria, Planctomycetes, Chloroflexi, and Verrucomicrobia decreased with depth. *Cytophaga aurantiaca* represented the most abundant permafrost layer contig, mapping 5.0 ± 4.0% of all permafrost counts. Bacteriodetes increased with depth, especially within TZ2 and PF. Here, one Sphingobacteriales contig alone constituted 9.1 ± 5.3% of all counts at 70−80 cm and mapped more than half of all counts in TZ2.

#### 3.3.2. Microeukaryotic permafrost community

Of the total eukaryotic community, 33.6% on average were made up of the supergroup Stramenopiles-Alveolata-Rhizaria (SAR) contigs (5.3 ± 3.7% of total counts), its relative abundance increasing in the transition zones (Fig. 3C). Eukaryotes were further represented by Opisthokonta (2.0 ± 3.8% of total counts), a clade that includes the animal and fungal kingdoms. Opisthokonta decreased with depth, although one insect rRNA contig created high counts in one 90 cm deep triplicate. Fungi were overall present at low relative abundance of 1.0 ± 1.1% of the total reads, representing 9.1% of the eukaryotic counts. The abundance of Archaeplastida counts was highest in the top 10 cm and in almost the complete transition zone at 50–90 cm (3.3 ± 1.3%) but had low relative abundances at 10–50 cm depth (<1%). Amoebozoa were mainly represented in the active layer and TZ1 (1.6 ± 0.5% as opposed to 0.9 ± 0.2% in TZ2 and PF).

#### 3.3.3. Bacteria-feeding prokaryote and protozoa community

Out of the 402 rRNA contigs assigned to predatory taxa, 258 were bacterial and 144 eukaryotic. The total proportion of predatory taxa within the total community ranged from 7.93–26.27%, peaking at 40–50 cm depth within the permafrost soil first thawed after the collapse. The ratio of protozoa to bacterial predators indicated a higher abundance of protozoa with depth, while predatory bacteria dominated especially within the active layer (Tab. 3). We tested if predator relative abundance significantly correlated with the total rRNA community composition. This revealed that both the abundance of bacterial predators (F.Model = 2.32, R^2^ = 0.225, p = 0.034) and protozoa (F.Model = 2.63, R^2^ = 0.247, p = 0.014) significantly shaped the prokaryotic, but not eukaryotic community (Tab. 2). Several non-predatory prokaryote abundances decreased in depths where predator abundance increased. Hence, we selected Alphaproteobacteria, Actinobacteria, and Bacteroidetes as potential prey and found that a similar significance level and effect size for these as for the total non-predatory community (Supp. Tab. 3). Some bacteria were significantly affected by protozoan predation (Sphingobacteriales F.Model = 4.59, R^2^ = 0.365, p = 0.004; Burkholderiales F.Model = 4, R^2^ = 0.333, p = 0.015, Supp. Tab. 3), while Nitrosomonadales was not.

**Tab. 3.** Average relative abundances in % per total prokaryotic community including prokaryotes (prok* = prokaryotes excluding bacterial predators) and their predators (= pred) per depth horizon. These include all predatory taxa (pred), the bacterial predatory (bacpred) orders Bdellovibrionales and Myxococcales, as well as eukaryotic protozoan superkingdoms (prot). Standard deviation is given in ±, accounting for the variation among triplicates per depth. Ratios of total predators to potential prey (prok*) and protozoan to bacterial predator ratio with depth.

| depth [cm] | 0-10 | 10-20 | 20-30 | 30-40 | 40-50 | 50-60 | 60-70 | 70-80 | 80-90 | 90-100 |
| --- | --- | --- | --- | --- | --- | --- | --- | --- | --- | --- |
| prok* | 79.29 ± 0.11 | 89.93 ± 0.01 | 87.25 ± 0.05 | 77.77 ± 0.04 | 71.25 ± 0.04 | 77.81 ± 0.09 | 82.35 ± 0.06 | 83.46 ± 0.02 | 78.04 ± 0.12 | 87.18 ± 0.09 |
| pred | 12.52 ± 0.14 | 7.93 ± 0.04 | 11.21 ± 0.07 | 20.16 ± 0.17 | 26.27 ± 0.33 | 19.44 ± 0.31 | 15.06 ± 0.16 | 12.68 ± 0.12 | 10.90 ± 0.10 | 8.56 ± 0.06 |
| bacpred | 7.56 ± 0.12 | 6.09 ± 0.04 | 8.85 ± 0.09 | 16.24 ± 0.21 | 15.31 ± 0.27 | 10.60 ± 0.33 | 4.72 ± 0.06 | 2.67 ± 0.04 | 1.28 ± 0.02 | 2.66 ± 0.03 |
| \| Bdellovibrionales | 0.14 ± 0.00 | 0.18 ± 0.00 | 0.24 ± 0.01 | 0.38 ± 0.01 | 0.53 ± 0.03 | 0.44 ± 0.05 | 0.67 ± 0.07 | 0.81 ± 0.04 | 0.55 ± 0.03 | 0.23 ± 0.01 |
| \| Myxococcales | 6.38 ± 0.10 | 5.91 ± 0.05 | 8.57 ± 0.09 | 15.38 ± 0.23 | 14.70 ± 0.30 | 10.11 ± 0.37 | 3.97 ± 0.06 | 1.83 ± 0.03 | 0.72 ± 0.01 | 2.40 ± 0.03 |
| \| \| Haliangiaceae | 3.23 ± 0.14 | 1.97 ± 0.03 | 2.73 ± 0.06 | 7.57 ± 0.21 | 8.81 ± 0.45 | 6.55 ± 0.59 | 0.80 ± 0.03 | 0.33 ± 0.01 | 0.11 ± 0.00 | 0.88 ± 0.02 |
| \| \| Blrii41 | 0.25 ± 0.05 | 0.77 ± 0.14 | 1.50 ± 0.32 | 3.55 ± 0.83 | 1.90 ± 0.36 | 0.77 ± 0.20 | 0.20 ± 0.04 | 0.06 ± 0.01 | 0.04 ± 0.01 | 0.25 ± 0.07 |
| prot | 4.97 ± 0.18 | 1.84 ± 0.02 | 2.36 ± 0.02 | 3.92 ± 0.05 | 10.96 ± 0.41 | 8.84 ± 0.27 | 10.34 ± 0.25 | 10.02 ± 0.19 | 9.62 ± 0.16 | 5.91 ± 0.09 |
| \| Amoebozoa | 2.26 ± 0.31 | 0.98 ± 0.03 | 0.87 ± 0.02 | 1.43 ± 0.05 | 1.91 ± 0.11 | 1.72 ± 0.11 | 1.70 ± 0.13 | 1.05 ± 0.05 | 0.91 ± 0.06 | 0.65 ± 0.03 |
| \| Excavata | 1.15 ± 0.37 | 0.08 ± 0.01 | 0.19 ± 0.01 | 0.37 ± 0.03 | 0.67 ± 0.07 | 0.46 ± 0.04 | 0.96 ± 0.10 | 0.92 ± 0.11 | 0.83 ± 0.11 | 0.54 ± 0.10 |
| \| SAR | 1.47 ± 0.04 | 0.72 ± 0.01 | 1.21 ± 0.02 | 1.99 ± 0.05 | 8.13 ± 0.51 | 6.49 ± 0.33 | 7.47 ± 0.30 | 7.84 ± 0.23 | 7.74 ± 0.20 | 4.64 ± 0.10 |
| \| \| Cercozoa | 0.99 ± 0.04 | 0.60 ± 0.01 | 0.58 ± 0.01 | 0.99 ± 0.01 | 2.56 ± 0.06 | 2.40 ± 0.05 | 3.24 ± 0.08 | 3.26 ± 0.09 | 3.13 ± 0.10 | 2.22 ± 0.06 |
| \| \| Ciliophora | 0.48 ± 0.02 | 0.12 ± 0.00 | 0.63 ± 0.03 | 1.00 ± 0.08 | 5.57 ± 0.87 | 4.09 ± 0.56 | 4.22 ± 0.50 | 4.59 ± 0.37 | 4.61 ± 0.30 | 2.42 ± 0.15 |
| \| Hacrobia | 0.01 ± 0.01 | 0.01 ± 0.00 | 0.01 ± 0.00 | 0.01 ± 0.00 | 0.03 ± 0.02 | 0.04 ± 0.04 | 0.07 ± 0.06 | 0.01 ± 0.00 | 0.00 ± 0.00 | 0.01 ± 0.01 |
| pred:prok* | 0.16 | 0.09 | 0.13 | 0.26 | 0.37 | 0.25 | 0.18 | 0.15 | 0.14 | 0.10 |
| protozoa:bacpred | 0.66 | 0.30 | 0.27 | 0.24 | 0.72 | 0.83 | 2.19 | 3.76 | 7.50 | 2.23 |

##### 3.3.3.1. Bacterial predators

Bacterial predatory rRNA contigs constituted on average 7.6% of the whole microbial community. They included the proteobacterial clades of Myxococcales, and in lower abundances Bdellovibrionales, as well as three *Lysobacter* rRNA contigs (Fig. 3B, Tab. 3). Myxococcales was the overall most abundant predator order with 205 rRNA contigs (7.1 ± 6.1% of total counts, 92.7% of bacterial, and 48.6% of all grazers).

##### 3.3.3.2. Protozoa

Protozoa included 144 rRNA contigs (6.9 ± 1.6% of total counts), mostly represented by rRNA contigs from the SAR clade. Cercozoa (2.0 ± 1.2%) and Ciliophora (2.8 ± 2.5%) dominated the protozoan community (Fig. 3D, Tab. 3). They were abundant in both transition zones and reached a maximum abundance at 60−70 cm depth (Fig. 3A), the deepest thaw level recorded in 2019 (Fig. 1D). Amoebozoa were less abundant (1.3 ± 0.7%), with *Lobosa* rRNA contigs being most abundant at 40–60 cm within the first thaw horizon TZ1 (Fig. 2D). Of SAR counts, 70.4% mapped in the transition zones, reflected by even representation of Alveolates and Rhizaria, while Stramenopiles mainly occurred in TZ2.

## 4. Discussion

In this study we aimed to elucidate the response of the total active permafrost microbial community in up to 26’500-year-old material under *in situ* abrupt thaw conditions. We successfully sequenced soil samples down to depths of ancient, intact permafrost and found a microbial bloom at recently thawed layers and changing abundances of microeukaryotes, including bacteria-feeding protozoa. Using metatranscriptomics, we were able to capture the rapid dynamics of fast-growing taxa, suggesting a microbial bloom upon thaw. To understand which taxa are responsible for the increased activity upon thaw, we here analysed the total community for prokaryotic and eukaryotic taxa associated with a potentially copiotrophic as well as bloom-controlling predatory lifestyle. In our study, the overall alpha diversity was similar to former metatranscriptomic research on permafrost microbiomes (Tveit *et al*., 2015), or even higher (Hultman *et al*., 2015, Müller *et al*., 2018, Schostag *et al*., 2019). The active layer and thawed permafrost indicated similar overall richness and abundance of rRNA contigs and hence potential microbial activity. Yet, the deepest thaw horizon indicated overall fewer and less active taxa than expected, compared to former amplicon-based findings (Scheel *et al*., 2022), although a few fast-growing taxa (Burkholderiales, Sphingobacteriales, Nitrosomonadales, and Myxococcales) dominated these depths. In this vulnerable but changing ecosystem, this highlights the importance of applying RNA-based techniques compared to DNA-based approaches in assessing each component of the total active community during environmental stress.

### 4.1. Community composition in eroding permafrost

The use of total RNA sequencing enabled the description of the entire soil community with full-length rRNA analysis, including prokaryotic and eukaryotic microbial taxa. The few metatranscriptomic studies from permafrost environments confirmed similar relative abundances of all domains (Hultman *et al*., 2015; Tveit *et al*., 2015; Schostag *et al*., 2019). While these studies have detected eukaryotic taxa, with 10.9% on average we found a higher abundance within the total active community (Tveit *et al*., 2015). The overall low fungal abundances in our study could relate to the rapid decrease of fungi upon thaw (Hultman *et al*., 2015). The ecological importance of bacterivorous microeukaryotes in temperate soils is a rising field of research, while poorly understood in permafrost.

In our study, most prokaryotes were represented by proteobacterial taxa, which were formerly reviewed as major taxon of both intact permafrost and active layer microbial communities (Jansson & Tas, 2014; Malard & Pearce, 2018). We noticed low relative abundance of Firmicutes rRNA and higher relative abundance of Planctomycetes than expected based reviewed DNA-based microbiomes (Jansson & Tas, 2014). This potentially indicates lower and higher activity, of Firmicutes and Planctomycetes, based on rRNA compared to amplicons. The relative abundance of Bacteroidetes increased with depth, indicating that some of its members are active in permafrost, as opposed to migrating from the active layer.

Generally, total RNA sequencing allows for insights into taxa that are active or at least alive at the timepoint of sampling, generally with a higher taxonomic resolution compared to DNA-based and particularly amplicon sequencing. However, we found that almost half of all rRNA reads could not be annotated. Taxonomic assignments of rRNA contigs decreased at taxonomic levels at higher resolution than class level. This might indicate that many of the taxa found within permafrost either remain new to common microbial database (SILVA version 138), or potentially include a high abundance of viral RNA (Wu *et al.,* 2022). Ultimately, this calls for further investigation of the so far undiscovered functional potential and microbial diversity within this dataset, including assembly of metagenome-assembled genomes (Sipes *et al*., 2021).

### 4.2. Thaw and other abiotic drivers

In this study, abiotic factors, such as the soil layer, soil organic matter content, moisture, and predation significantly shaped the overall microbial community. Yet only the active layer and permafrost communities significantly differed from each other. Prokaryotic trends reflected those of the entire community, while changes in the eukaryotic community were furthermore significantly described by pH and indicated significantly different communities in each layer. This pattern can perhaps be explained with higher mobility in eukaryotic taxa that allows migration to the optimum habitat while other immobile microbial taxa cannot – especially when thaw decreases the former dispersal limits for microbial taxa, that depend on soil moisture for motility (Bottos *et al*., 2018).

Many microbial taxa, including predatory taxa, responded rapidly to thaw. After the former dispersal-limiting ice lens melted, we expect a slow and gradual migration from the active to deeper thawed layers. This could explain, why no significant differences in rRNA community composition were found between the two thawed depths. This supports the finding that the most recently thawed soil (TZ2) did not contain a higher relative abundance of rRNA for permafrost endemic taxa than TZ1. A bloom of such taxa by reactivating spore could be dilute by the effect of migration over time.

The strong correlation between soil moisture and the microbial community can be an artifact as the highest soil moisture values coincide with depths that are also assigned as active layer. This might be why soil moisture, SOM and layer significantly correlated with each other (ANOVA p <0.005; Supp. Tab. 2). Still, these edaphic properties depict the rather short-term character of the site. And variance partitioning showed that even a combination of short-, long-term and predatory effects could not explain the majority of variance in community composition.

### 4.3. Copiotrophs in thawing permafrost soil

Within the prokaryotic community, the most abundant taxa overall also dominated within the recently thawed permafrost material (Supp. Fig. 3), in agreement with our former DNA amplicon-based study in this site (Scheel *et al*., 2022). Many of these are conditionally rare, but metabolically important autotrophic taxa, that can respond rapidly to changing environmental factors (Shade *et al*., 2014). Overall, proteobacterial contigs were the most abundant and encompassed the most phylogenetically diverse groups. Especially Betaproteobacteria made up 8 of the 20 most abundant taxa (all Gallionellaceae) and 1/4 of all taxa in both thawed layers. Betaproteobacteria include many copiotrophic representatives, that were formerly observed to quickly respond to freshly available resources upon permafrost thaw (Hurst, 2019, Schostag *et al*., 2019). Alpha- and Betaproteobacteria include many copiotrophic representatives that can quickly adapt to changing environmental conditions (Schostag *et al*., 2019). Especially short-term stressors were seen to trigger a dominance of soil copiotrophs (Koyama *et al*., 2014, Bang-Andreasen *et al*., 2020).

The thaw of permafrost soils poses a large potential liberation of organic and inorganic nitrogen into the arctic subsoils for microbial uptake (Keuper *et al*., 2012, Voigt *et al*., 2017, Voigt *et al*., 2020, Marushchak *et al*., 2021, Wegner *et al*., 2022). Here we show a large potential for rapid N-cycling when permafrost thaws. Notably, most contigs with maximum relative abundances at this depth (70 cm), were the abundant Gallionellaceae (Nitrosomonadales, Betaproteobacteria) (Hallbeck & Pedersen, 2014), which contain nitrate-dependent iron-oxidising taxa (He *et al*., 2016). This family was previously found in intact ancient permafrost (Alawi *et al*., 2007, Müller *et al*., 2018, Scheel *et al*., 2022). Nitrosomonadales occurred at the first erosion year’s maximum thaw depth of 70 cm, which is associated with the abundance of nitrogen pools in permafrost soils (Schostag *et al*., 2019). As nitrifier, this functional guild might supply the substrate needed for consequent denitrification processes and hence release of N_2_O from thawing permafrost. This emission has been found to place thawing permafrost as top N_2_O source beside tropical rainforests (Voigt *et al*., 2017), likely due to the 7 times higher N content and elevated mineralisation rates at the permafrost Tab. compared to the rooting zone (Keuper *et al*., 2012). Although nutrient measurements and nitrification gene abundance are needed to confirm active gene expression, our results confirm the ecological importance of nitrogen cycling in thawing permafrost, as suggested before (Keuper *et al*., 2020, Monteux *et al*., 2020).

Another Betaproteobacteria major order, Burkholderiales, increased steadily towards the second, deepest thaw depth. Especially Comamonadaceae, a family that consists of many copiotrophic spore-formers, rapidly respond to fresh input of labile nutrients (Fierer *et al*., 2007, Ho *et al*., 2017). This suggests that these mineral-weathering Burkholderiales (Naylor *et al*., 2022) might be readily responsive to thawing and hence bioavailable permafrost carbon.

Similarly, the particularly active Bacteroidetes order Sphingobacteriales dominated recently thawed permafrost, as found to be connected to thaw before (Müller *et al*., 2018). In our study site, Sphingobacteriales had the highest relative abundance within the total amplicon-based community composition (Scheel *et al*., 2022). They then constituted more than half of all counts between 70−90 cm depth in 2020. This stands in contrast with very low 16S rRNA-based abundances one year earlier (Scheel *et al*., 2022), which could mean that Sphingobacteriales particularly responds rapidly within recently thawed permafrost. Several Bacteriodetes taxa were documented as initial metabolisers of labile carbon (Fierer *et al*., 2007, Ho *et al*., 2017, Stone *et al*., 2023), which highlights their potential impact on thawed permafrost carbon fluxes. We confirm Bacteroidetes as a major component of the microbial response to thaw (Frank-Fahle *et al*., 2014, Coolen & Orsi, 2015, Deng *et al*., 2015, Burkert *et al*., 2019). Bacteroidetes has been previously found to be abundant especially at the upper permafrost limit (Müller *et al*., 2018, Tripathi *et al*., 2018).

### 4.4. Predation as response to thaw dynamics and biotic driver

Bacterial and protozoan predators significantly impacted the overall non-predatory prokaryotes and showed strong correlation with several orders typically representing fast-growing bacterial lifestyles, such as Sphingobacteriales. In a former study investigating the impact of nematodes and protozoa on different bacterial lifestyles under nutrient addition showed that oligotrophic taxa often served as prey, the overall bacterial growth being more impacted by nutrient availability than predation (Zelenev *et al*., 2006). This underlines our findings that predator abundance was important to the system as biological control of microbial blooms but did not act as the main driver of taxonomic changes across depths. This further highlights the lack of knowledge in respect to ecological diversity and function of protists in soil, but especially permafrost systems (Geisen & Bonkowski, 2018).

#### 4.4.1. Overall potential prey and predation distribution

Several microorganisms have protective measures against predation (Trap *et al*., 2016). These include especially gram-positive bacteria, such as bacilli and cocci, as well as Actinobacteria and Firmicutes, which are especially abundant in permafrost (Trap *et al*., 2016). This protection can be achieved by the formation of filaments, biofilms and colonies, cell size, shape, pigments, and toxins. Here, we noted the decrease of Actinobacteria in the first transition zone, where predators were abundant. Although protozoan abundance correlated with Actinobacteria abundance, this finding seems contradictory, as representatives of this phylum can form filaments. This could indicate that other environmental processes override the potential impact of predation in our system (Fig. 2). In contrast, gram-negative bacteria are suitable prey especially to myxobacteria and protists (Trap *et al*., 2016). Hence, the bloom of gram-negative Proteobacteria in the thaw zone led us to the idea that potentially fast-growing prey support a consecutive predator succession. Myxobacteria were found to select on gram-negative bacteria, while protozoa showed less selective feeding (Zhang & Lueders, 2017). This might be reflected in the overall stronger correlation of protozoan abundance with the total and sub-communities. Although species specific feeding had been found among protozoa (Pedersen *et al*., 2011, Rønn *et al*., 2001), we could not clearly confirm prey taxa, nor taxa that potentially benefited from predation in the scope of this study.

#### 4.4.2. Bacterial predators dominate shallow thaw

We found that Myxococcales was the most abundant order after Nitrosomonadales, with abundances in the range of former findings from Arctic soils (Malard & Pearce, 2018). Bacterial predators and particularly Myxococcales play key roles in microbial food webs, as recently demonstrated (Dai *et al*., 2021, Petters *et al*., 2021). Former studies have shown Myxococcales to be abundant in active layer samples (Inglese *et al*., 2018, Malard & Pearce, 2018, Romanowicz & Kling, 2022). Within the active layer, seasonal prokaryotic blooms could support higher predator abundances. We hypothesise that the formation of spore-containing fruiting bodies in Myxococcales (Huntley *et al*., 2011) offers resistance to the winter freeze-off, contrary to the freeze-induced mortality of prey taxa, might supply the growth substrate for Myxococcales in spring (Bang-Andreasen *et al*., 2020). Bdellovibrionales were found in the deeper thawed layers. These flagellated bacteria have a lower prey range and potentially thrive were competition with Myxococcales is lower (Petters *et al*., 2021). With overall 0.4% of all rRNA contigs, we found a similar relative abundance of predatory bacteria as formerly observed in a peatland (Petters *et al*., 2021).

#### 4.4.3. Protozoa as fast responding predators

Protozoa taxa included Cercozoa, Amoebozoa, and Ciliophora in agreement with former studies (Schostag *et al*., 2019; Shatilovich *et al*., 2009). Especially Ciliophora and Cercozoa responded to permafrost thaw and reached maximum abundances at 70 cm depth, the deepest thaw depths in the first erosion year. Protozoa were most abundant in both thawed permafrost layers, making them a more abundant predator than Myxobacteria, which in turn dominated the active and shallow first thaw layer (Fig. 3B & D). One potential explanation for the high abundance of protozoa in freshly thawed permafrost could be that several cyst- and resting-stage-forming protist taxa can respond quickly to environmental changes (Rønn *et al*., 2012).

Interestingly, among the few permafrost metatranscriptomic studies performed, increases in relative abundance of protozoan taxa with increasing temperature were not mentioned (Schostag et al, 2019), while an increase of Cercozoa was observed (Tveit et al, 2015). Although our relative abundances of Cercozoa matches those of former studies (Geisen *et al*., 2015, Tveit *et al*., 2015, Schostag *et al*., 2019), we expected higher overall abundances at the most recently available thaw horizon at 90 cm. Due to their relatively smaller size, Cercozoa reproduce faster than larger microeukaryotes (Rønn *et al*., 2012). Furthermore, members of this group have been shown to respond to permafrost thaw within days (Schostag *et al*., 2019), as well as to temperature increases in Arctic peat (Tveit *et al*., 2015). Ciliates and Amoeba respond very rapidly to changes in soil moisture (Coleman & Wall, 2015) and temperature (Schostag *et al*., 2019) and have even been found in ancient Siberian permafrost (Shatilovich *et al*., 2009).

Even though a faster response is more likely in prokaryotic predators, due to longer generation times of protozoans (Ekelund *et al*., 2002). Our findings suggest that more mobile predators dominate during the initial stages of the thaw-triggered microbial bloom, while during later stages of thaw increased relative abundance of the less mobile bacterial predators occurred. Furthermore, protozoans might also benefit from the unselective feeding mechanisms in contrast to the more selective Myxobacteria (Zhang & Lueders, 2017). This potentially explains why we observed changes in distribution with depth and significant impact of protozoan, but not myxobacterial abundance on potential prey taxa.

### 4.5. Microbial bloom during thaw and potential coalescence

Predation bears the potential to reduce and hence control microbial population growth, but also has been found to lead to higher yields in more diverse artificial incubation communities, likely due to reduced competition from prey taxa (Saleem *et al*., 2012). We noted that both prokaryotic and eukaryotic taxa contributed to higher activity in the thawed layers. Thus, our findings of higher RNA:DNA ratios at the freshly thawed 50 and 80 cm depths indicated higher ribosomal relative abundance of both domains at these depths (Tab. 1, Supp. Fig. 1).

The difference in the composition of the active microbial community between more and less recently thawed permafrost layers could result from multiple ecological processes that simultaneously influence the microbial community composition. For example, 1) a steady migration of microorganisms from the active layer to the thawed layers, compared to the below permafrost layer, and 2) rapid microbial blooms in recently thawed, former permafrost layers. Ecological responses of soil microbiomes to environmental stress are non-linear and complex, arising from simultaneous stochastic and deterministic environmental selection (Doherty *et al*., 2020, Ernakovich *et al*., 2022). Even with low increase of absolute temperature, not only does thaw impose enormous stress on the microbiome, but Wang and colleagues found several species especially with the phyla of Proteobacteria and the order Sphingobacteriales to respond even to changes at low temperatures of 5–15 °C, as opposed to most temperate taxa at 15–25 °C (Wang *et al*., 2021).

A rapid bloom in ancient, former frozen permafrost, could stem from endemic spores and resting stages. Situated by the Zackenberg river, the oldest sand and rubble deposits (AO) indicate a fluvial origin, potentially as part of the Zackenberg river or glacial meltwater delta into the fjord roughly 30’000 years ago (Gilbert *et al*., 2017), while younger material could resemble a former pond, that later became overgrown (AM). Arctic fluvial habitats showed higher abundances of Bacteroidetes and Proteobacteria in a summer glacial river, but also the Arctic Lake Hazen sediments (Cavaco *et al*., 2019) and subglacial habitats (Achberger *et al*., 2017), which could explain their origin even in our samples of ancient and still intact permafrost. Simultaneously, dominance of Bacteroidetes were also found in permafrost metatranscriptomes during thaw (Hultman *et al*., 2015, Schostag *et al*., 2019).

In contrast to the idea of increased activity in former cold-adapted taxa, earlier findings showed the potential for coalescence of taxa from the active layer to thawing permafrost (Rillig *et al*., 2015). This process was formerly observed upon transplantation of grassland soil microbiomes onto permafrost (Monteux *et al*., 2020). Such mixing conditions occurs naturally during cryoturbation of permafrost, such as during a collapse (Gittel *et al*., 2014, Schnecker *et al*., 2014). In their previous incubation study, Monteux and colleagues found an increase of Bacteroidetes abundance upon thaw, after inoculating permafrost with grassland soil (Monteux *et al*., 2020). Both the processes of a thaw control as well as implantation of temperate soil have resulted in elevated nitrification and CO_2_ emissions (Monteux *et al*., 2020). This highlights the importance of understanding community dynamics in response to thaw. Older permafrost soils are now increasingly recognised as functionally limited due to thermally restricted conditions (Monteux *et al*., 2020, Wegner *et al*., 2022). Although we could not statistically separate the potential effects of isolated permafrost thaw and coalescence with the active layer community, they might supply a higher functional diversity to overcome this limitation and hence enhance carbon and nitrogen from thawed permafrost.

### 4.6. Conclusion

We have described the total prokaryotic and microeukaryotic community composition abrupt permafrost erosion stress in High Arctic Greenland based on total RNA sequencing of *in situ* samples. The composition of the total community was driven both by the abiotic factors related to the thawing state of the soil layer, soil moisture, and organic matter content. The microbial bloom we found within the transition zone, as indicated by large increases in the RNA:DNA ratio, enabled a diverse and active microbial food web. An increase of copiotrophic taxa in the freshly thawed permafrost, included a dominance of potentially fast-growing and -metabolising order Nitrosomonadales, Sphingobacteriales, and Burkholderiales. This bloom at thawed depths also formed the base of a solid bacteria-feeding community dominated by protozoan bacterivores in recently thawed soil and bacterial predators just below the historical active layer boundary. We found that predation additionally to abiotic processes, impacted the active microbial community composition and highlight the importance of trophic interactions as ecological response to permafrost collapse.

## 6. Acknowledgements

We thank the Zackenberg Research Station staff, the Greenland Ecosystem Monitoring program facilities and its field assistants for smooth sample handling of the sampling and samples.

## 7. Data availability statement

The raw sequence data of this study were deposited in the NCBI Sequence Read Archive and can be accessed through accession number PRJNA939404.

## Supplementary Material

**Supp. Fig. 1.**
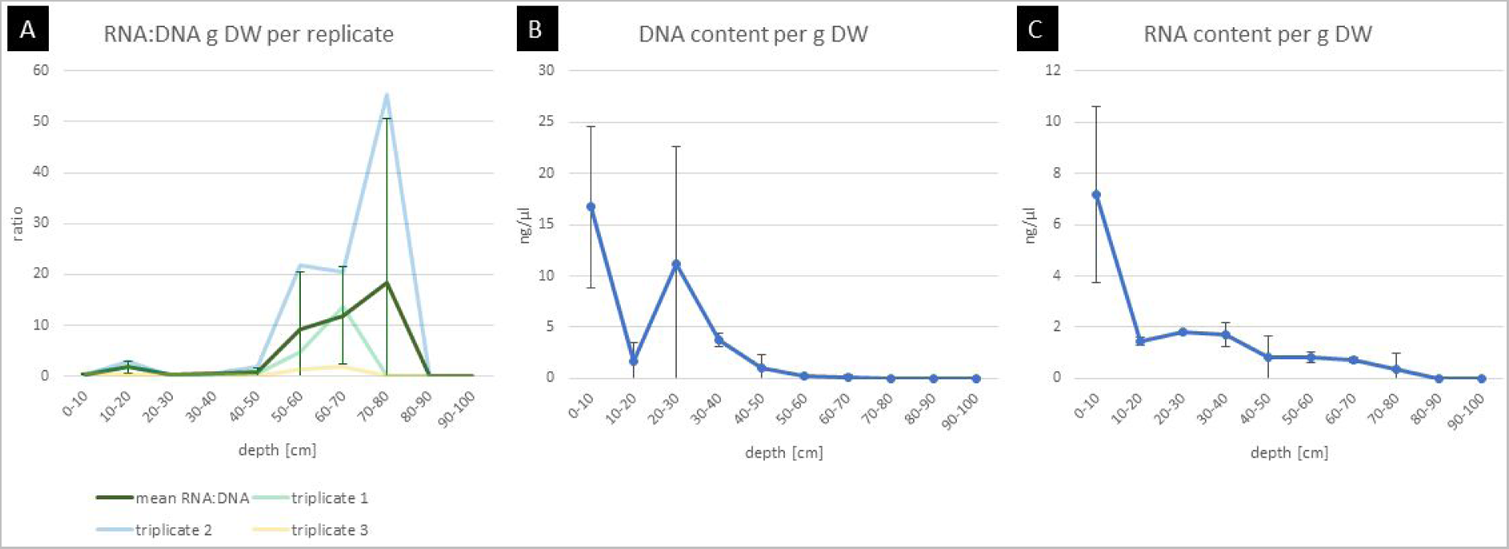
Dry weight (DW) normalised RNA:DNA ratio per gram sample was indicated for each extraction triplicate and the resulting mean. The mean co-extracted DNA (B) and RNA (C) content in ng/µl per g DW per depth as measured using TapeStation assays. Whiskers indicate the standard deviation among the extraction triplicates.

**Supp. Fig. 2.**
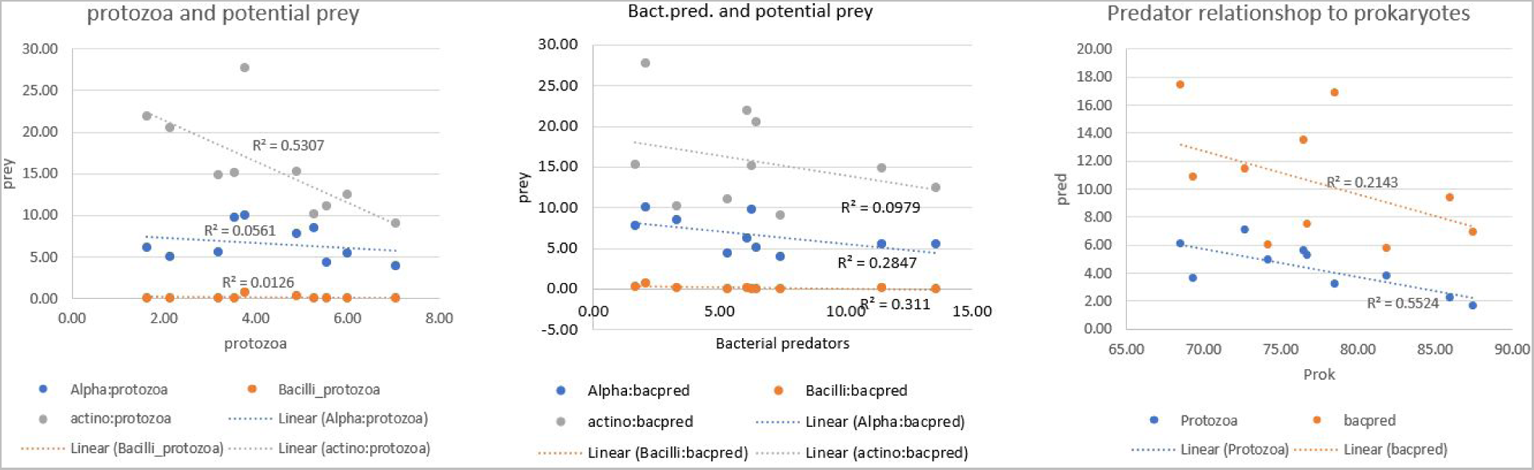
Testing potential correlation between predator and prey relative abundance per sample for selected taxa, including Bacilli, Alphaproteobacteria and Actinobacteria as prey. Protozoa (left), to bacterial predators (bacpred, middle) and the overall correlation between non predatory prokaryotes (prok) to protozoa and bacterial predators (right). The linear fit is given with an R^2^ each.

**Supp. Fig. 3.**
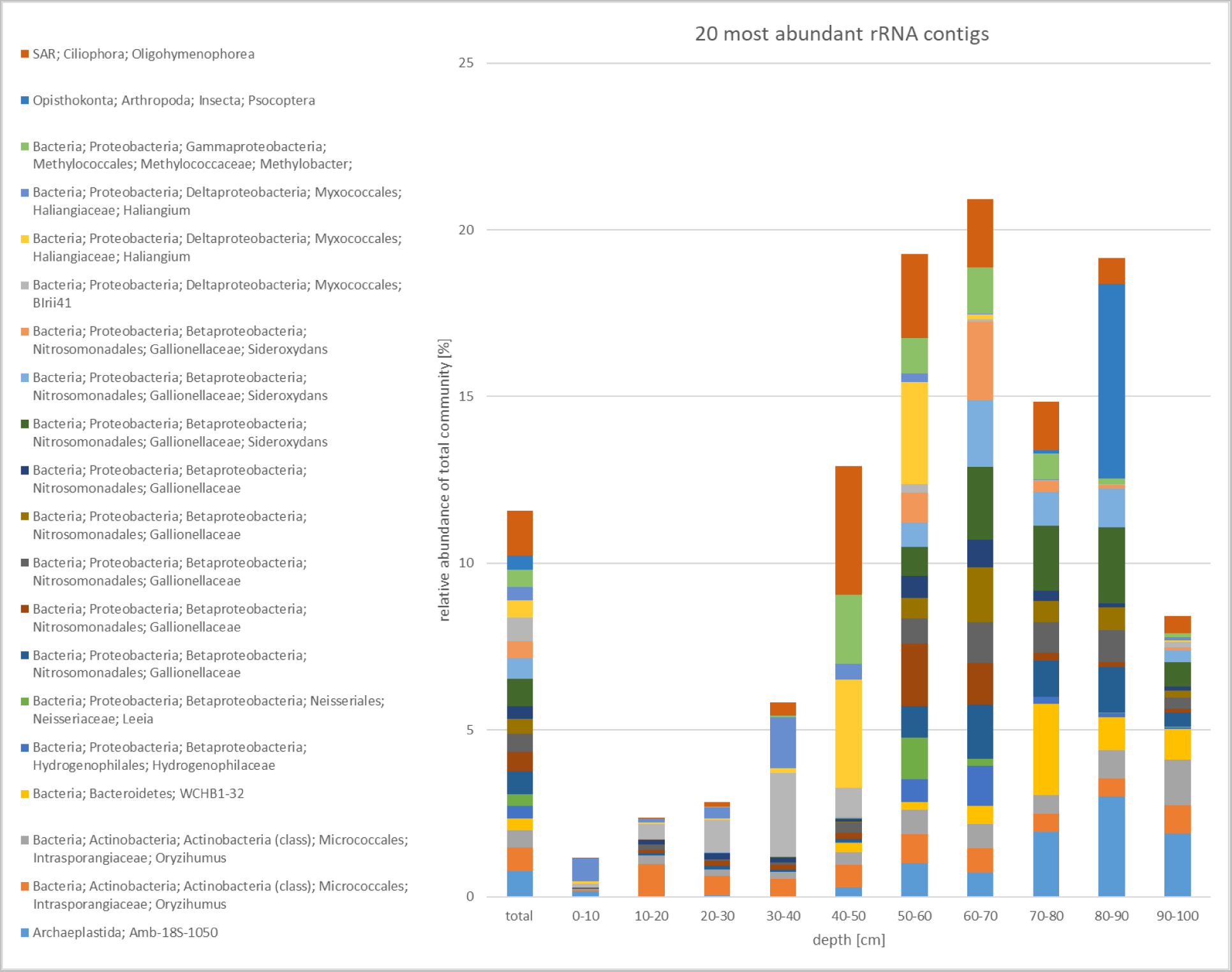
The 20 rRNA contigs with highest total relative abundance in the overall annotated community composition are given in relative abundance per 10 cm depth interval across all domains. Taxonomic annotation is given in Domain; Kingdom; Phylum; Class; Order; Family; Genus; Species.

**Supp. Tab. 1.** The partition of SSU-annotated reads (rRNA) and neither SSU-nor LSU-annotated (mRNA) reads of the total reads. Diversity measures here include the Shannon index (S) and number of rRNA contigs mapped per depth. The total number included all rRNA contigs, excluding these annotated as “Chloroplast” or taxonomic levels starting with “Unknown”. Standard deviation from triplicates is given as ±.

| depth [cm] | rRNA % | RIN | mRNA % | S | richness |
| --- | --- | --- | --- | --- | --- |
| 0-10 | 77.39 ± 1.08 | 0.4 ± 0.7 | 7.60 ± 2.73 | 7.44 ± 0.10 | 3727 ± 1 |
| 10-20 | 67.40 ± 8.90 | 1.2 ± 1.2 | 18.69 ± 0.52 | 7.15 ± 0.03 | 3727 ± 2 |
| 20-30 | 72.67 ± 3.99 | 0.5 ± 0.8 | 14.50 ± 3.09 | 7.17 ± 0.02 | 3724 ± 4 |
| 30-40 | 66.85 ± 3.42 | 0.7 ± 1.2 | 8.44 ± 3.48 | 7.19 ± 0.02 | 3727 ± 1 |
| 40-50 | 77.65 ± 4.82 | 1.3 ± 0.4 | 43.13 ± 56.71 | 6.68 ± 0.02 | 3719 ± 8 |
| 50-60 | 75.74 ± 1.08 | 1.3 ± 0.5 | 9.51 ± 1.25 | 6.22 ± 0.20 | 3720 ± 11 |
| 60-70 | 73.76 ± 7.28 | 1.6 ± 0.5 | 7.79 ± 1.30 | 6.20 ± 0.31 | 3718 ± 12 |
| 70-80 | 80.86 ± 2.43 | 1.8 ± 0.2 | 6.15 ± 1.70 | 6.20 ± 0.33 | 3376 ± 500 |
| 80-90 | 75.14 ± 6.49 | 1.1 ± 1 | 12.70 ± 7.57 | 5.83 ± 0.35 | 3707 ± 3 |
| 90-100 | 55.39 ± 20.91 | 0 ± 0 | 23.38 ± 8.18 | 6.04 ± 0.89 | 3394 ± 477 |

**Supp. Tab. 2.** Results given as R^2^ value for permutational multivariate analysis for the biotic and abiotic soil properties against each other (age categories, layer, soil organic matter (SOM) and moisture (H2O), pH, bacterial predator (bacpred), and protozoa relative abundance. P-values < 0.05 were considered significant with * p<0.05, ** p<0.01, *** p<0.005. Non-significant results were summarised as NS.

|  | age | layer | SOM | H<sub>2</sub>O | pH | bacpred | protozoa |
| --- | --- | --- | --- | --- | --- | --- | --- |
| depth | 0.368* | 0.518** | 0.391* | **0.63\*\*\*** | 0.409* | NS | 0.353* |
| age | - | 0.406* | NS | NS | NS | NS | 0.493* |
| layer | - | - | **0.699\*\*\*** | **0.787\*\*\*** | NS | NS | 0.705** |
| SOM | - | - | - | **0.768\*\*\*** | 0.463* | 0.409* | 0.394* |
| H<sub>2</sub>O | - | - | - | - | **0.619\*\*\*** | NS | 0.473* |
| pH | - | - | - | - | - | NS | NS |
| bacpred | - | - | - | - | - | - | NS |

**Supp. Tab. 3.** Results of permutational multivariate analysis on the mean Bray-Curtis dissimilarities per depths between potential prey taxa (Alphaproteobacteria, Actinobacteria, Bacteroidetes) and potentially copiotrophic taxa (Burkholderiales, Sphingobacteriales). These were tested against the age and layer categories, and predator abundance. P-values < 0.05 were considered significant with * p<0.05, ** p<0.01, *** p<0.005. Non-significant categories, pair-wise contrasts, and taxa were removed, including Nitrosomonadales, bacterial predator abundances and pairwise contrasts of age categories.

|  | Potential prey taxa |  |  |  |  |  |  |  |  | Potentially copiotrophic taxa |  |  |  |  |  |
| --- | --- | --- | --- | --- | --- | --- | --- | --- | --- | --- | --- | --- | --- | --- | --- |
|  | α-Proteo. |  |  | Actinobact. |  |  | Bacteriod. |  |  | Burkhold. |  |  | Sphingobact. |  |  |
| Parameter | F | R<sup>2</sup> | p | F | R<sup>2</sup> | p | F | R<sup>2</sup> | p | F | R<sup>2</sup> | p | F | R<sup>2</sup> | p |
| AL-PF |  |  | *** |  |  | * |  |  | * |  |  |  |  |  | * |
| TZ1-PF |  |  | * |  |  |  |  |  |  |  |  |  |  |  |  |
| TZ2-PF |  |  | * |  |  |  |  |  |  |  |  |  |  |  |  |
| protozoa | 2.66 | 0.25 | * | 2.53 | 0.24 | * | 3.32 | 0.294 | * | 4 | 0.333 | * | 4.59 | 0.365 | *** |

